# Chronic Ethanol Differentially Modulates Glutamate Release from Dorsal and Ventral Prefrontal Cortical Inputs onto Rat Basolateral Amygdala Principal Neurons

**DOI:** 10.1101/558189

**Authors:** Molly M. McGinnis, Brian C. Parrish, Ann M. Chappell, Brian A. McCool

**Affiliations:** Department of Physiology and Pharmacology, Wake Forest School of Medicine, Winston-Salem, NC 27157

## Abstract

The medial prefrontal cortex (mPFC) and the basolateral amygdala (BLA) have strong reciprocal connectivity. Projections from the BLA to the mPFC can bidirectionally modulate anxiety-related behaviors but it is unclear if the same is true for mPFC to BLA projections. Our laboratory is specifically interested in withdrawal-related anxiety-like behavior and the underlying synaptic plasticity. Here, we use optogenetics and chemogenetics to characterize the neurophysiological and behavioral alterations produced by chronic ethanol exposure and withdrawal on dorsal mPFC/prelimbic (dmPFC/PL) and ventral mPFC (vmPFC/IL) terminals in the BLA. We exposed adult male Sprague-Dawley rats to chronic intermittent ethanol (CIE) using vapor chambers, measured anxiety-like behavior on the elevated zero maze (EZM), and used electrophysiology to record glutamatergic and GABAergic responses in BLA principal neurons. We found that 24-hour withdrawal following a 7-day CIE exposure significantly increased the glutamate release probability from PL/dmPFC terminals, but significantly decreases the glutamate release probability from IL/vmPFC terminals. Chemogenetic inhibition of PL/dmPFC terminals in the BLA attenuated the increased withdrawal-dependent, anxiety-like behavior. These data demonstrate that chronic ethanol exposure and withdrawal strengthens the PL/dmPFC – BLA pathway but weakens the IL/vmPFC – BLA pathway. Moreover, we provide novel evidence that the PL/dmPFC – BLA pathway can modulate anxiety-like behavior. These findings suggest that mPFC-BLA circuits known to regulate the acquisition of aversive behaviors are up-regulated by chronic ethanol while those involved with the extinction of these behaviors are down-regulated.

**Significance Statement:** Accumulating evidence suggests that the medial prefrontal cortex and its projections to the basolateral amygdala bidirectionally modulate fear-related behaviors. Since the neuronal circuits for fear and anxiety are thought to overlap, we sought to examine the role of prelimbic and infralimbic subdivisions of the medial prefrontal cortex and their inputs to the basolateral amygdala in regulating anxiety. Specifically, we focused on alcohol withdrawal-induced anxiety-like behavior, which is a commonly reported cause of relapse in human alcoholics. In our study, we used optogenetics and chemogenetics to demonstrate, for the first time, that withdrawal from chronic ethanol exposure strengthens prelimbic synapses, but weakens infralimbic synapses in the basolateral amygdala and that inhibiting glutamate release from prelimbic terminal in the basolateral amygdala reduces anxiety-like behavior.

## Introduction

Over the past few decades, epidemiological and clinical studies have highlighted the relationship between alcohol use disorder (AUD) and anxiety disorders (Smith and Randall, 2012). Comorbidity of these disorders is highly prevalent and is associated with a complex clinical presentation, which makes diagnosis and treatment challenging. Anxiety disorders in individuals with AUD is largely thought to be an artifact of alcohol withdrawal and can contribute to the maintenance of and relapse to pathological alcohol use (Kushner et al., 2000; Schuckit and Hesselbrock, 1994). The standard pharmacotherapy and psychotherapy protocols for AUD and anxiety treatments have had limited success, with roughly 80% of individuals relapsing following treatment (Driessen et al., 2001; Kushner et al., 2005). A crucial step in the development of more effective treatments for AUD and co-occurring anxiety disorders is to identify the brain circuits altered by alcohol dependence that are critically involved in the regulation of emotion.

The basolateral amygdala (BLA) has long been established as an integral part of the neural circuitry that regulates emotion, including fear and anxiety (Davis, 1992; Gallagher and Chiba, 1996; Tovote et al., 2015). Inputs to the BLA are widespread and arise from both cortical and subcortical regions conveying sensory information from all modalities (McDonald, 1998). Afferent sensory information is processed locally within the BLA and is then projected to efferent brain regions that produce a variety of behavioral responses (Sah et al., 2003). Studies on the morphological and physiological properties of cells within the BLA have described two main types of neurons. Glutamatergic pyramidal neurons comprise approximately 85% of the cell population and are the primary projection neurons of the BLA (Mcdonald, 1982; McDonald, 1992; Washburn and Moises, 1992). The remaining cell population consists of GABAergic neurons, which are local circuit interneurons that exert tight inhibitory control over the pyramidal neurons (Carlsen, 1988; Muller et al., 2006). Thus, the generation of emotional responses is predominantly determined by the balance of excitatory and inhibitory neurotransmission within the BLA (Davis et al., 1994; McDonald and Betette, 2001). Importantly, both the anatomy and function of the BLA is highly conserved across species (Janak and Tye, 2015), allowing for the use of translational animal models to study human conditions.

A leading neurobiological model of emotional regulation, supported by both animal and human data, is that the prefrontal cortex mediates the expression of negative affect through powerful top-down control of the amygdala (Phelps and LeDoux, 2005). Specifically, the dorsal and ventral subdivisions of the medial prefrontal cortex (mPFC) have strong reciprocal connectivity with neurons in the BLA (Little and Carter, 2013; McGarry and Carter, 2017) and have been implicated in controlling a variety of emotional-related behaviors (Arruda-Carvalho and Clem, 2015; Sotres-Bayon and Quirk, 2010). Interestingly, these different subregions and their projections to the BLA play dichotomous roles in fear learning and extinction. For example, the prelimbic or dorsal medial prefrontal cortex (PL/dmPFC) is involved in the acquisition of conditioned fear and fear expression (Arruda-Carvalho and Clem, 2014; Senn et al., 2014; Sierra-Mercado et al., 2011). Conversely, the infralimbic or ventral medial prefrontal cortex (IL/vmPFC) is critical for extinction learning and the inhibition of fear responses (Bloodgood et al., 2017; Bukalo et al., 2015; Senn et al., 2014; Sierra-Mercado et al., 2011). The ability of these brain regions to bidirectionally modulate such behaviors is likely due to the fact that inputs from the PL/dmPFC and IL/vmPFC make direct monosynaptic excitatory synapses onto projection neurons in the BLA, as well as interacting with local inhibitory circuits (Cho et al., 2013; Hübner et al., 2014; Likhtik, 2005; Strobel et al., 2015). Taken together, these data provide extensive evidence that the PL/dmPFC and IL/vmPFC projections to the BLA are important pathways for the regulation of fear. However, the role of these circuits is regulating other emotional behaviors, such as anxiety, has not been studied. Additionally, how these circuits might be altered under pathological conditions, such as during alcohol withdrawal, remains to be elucidated.

Evidence suggests that the key mechanisms underlying emotional behaviors such as fear and anxiety are similar in both animals and humans, and that these processes are likely mediated by overlapping neuronal circuits. Therefore, we hypothesized that projections from the PL/dmPFC and IL/vmPFC to the BLA would be altered by chronic ethanol exposure and may regulate alcohol withdrawal-induced anxiety-like behavior in an animal model. A commonly used method for including dependence in animal models of AUD is chronic intermittent ethanol (CIE) exposure. Using this model, our laboratory has demonstrated that withdrawal from CIE produces an increase in anxiety-like behavior (Morales et al., 2015), which is a result of increased presynaptic glutamate release onto BLA pyramidal neurons (Christian et al., 2013; Läck et al., 2007; Morales et al., 2018). These data suggest that glutamate inputs, from regions such as the mPFC, are likely altered by chronic ethanol exposure and withdrawal. Additionally, CIE impairs fear extinction and remodels glutamatergic mPFC neurons (Holmes et al., 2012) providing further evidence that specific mPFC projections to the BLA may be altered by chronic ethanol exposure and mediate withdrawal-induced anxiety. Here, we took a circuit-based approach using electrophysiology in combination with optogenetics and chemogenetics to examine PL/dmPFC and IL/vmPFC terminals in the BLA, the effects of CIE exposure, and their role in regulating anxiety-like behavior during withdrawal.

## Materials and Methods

### Animals

Male Sprague-Dawley rats were obtained from Envigo (Indianapolis, IN). Upon arrival, rats were pair-housed, given food and water *ad libitum*, and maintained on a reverse 12:12h light-dark cycle (lights on at 9 PM). Rats were aged ~5 weeks at the time of surgical manipulation and ~10 weeks at the time of behavioral manipulations/electrophysiology recordings. All animal care procedures were in accordance with the NIH Guide for the Care and Use of Laboratory Animals and experimental procedures were approved in advance by the Institutional Animal Care and Use Committee at Wake Forest.

### Stereotaxic Surgery

#### Viral Microinjection

Rats (n= 115; ~100g) were induced using 3-5% isoflurane anesthesia and maintained under continuous 1-3% isoflurane for the duration of the procedure with the oxygen flowmeter set to 1.0 L/min. Bilateral microinjections (1μL/side) of adeno-associated viral vectors containing channelrhodopin (AAV5-CamKIIα-hChR2(H134R)-EYFP; UNC Vector Core), Gi-coupled DREADD (AAV5-CamKIIα-hM4D(Gi)-mCherry; Addgene), or a control fluorophore (AAV5-CamKIIα-mCherry; UNC Vector Core) were delivered at a rate of 0.1μL/min over 10 min using a Harvard Apparatus pump (Holliston, MA). Injectors were left in place for an additional 5 min to allow for virus to diffuse. The medial prefrontal cortex was targeted using a Neurostar StereoDrive (Germany) with the following coordinates (in mm): dmPFC (3.20 AP, ± 0.50 ML, 2.75 DV) or vmPFC (3.20 AP, ± 0.50 ML, 4.75 DV) relative to bregma. Targeting the upper portion of the dmPFC and lower portion of the vmPFC allowed for maximal spatial segregation between subregions. At the end of surgery rats were given 3 mg/kg ketoprofen (Patterson Veterinary Supply, Charlotte, NC) for pain management and 2 mL warmed sterile saline. Sutures were removed 1 week later, and animals were pair-housed. Rats were allowed a total of 4 weeks to recover and for virus to express before further experimentation. Injection sites were confirmed by collecting coronal slices of the mPFC and visualizing EYFP and mCherry using fluorescence microscopy post-mortem. Rats with unintended viral spread were excluded.

#### Cannulation Surgery

For the behavioral studies using BLA microinjections, animals (n= 48; ~300g) underwent a second surgery 4 weeks after virus injection to implant bilateral guide cannulae using previously published methods (Diaz et al., 2011a; Läck et al., 2007). Briefly, rats were deeply anesthetized with pentobarbital sodium (50 mg/kg, IP) and stainless-steel guide cannulae (26 gauge; Plastics One, Roanoke, VA) of sufficient length were implanted to terminate 1 mm above the BLA using the following stereotaxic coordinates: −2.80 AP, ± 5.00 ML, 7.80 DV. Dental cement (Lang Dental Manufacturing Co., Wheeling, IL) was used to affix and stabilize the guide cannulae to the skull. Obturators (Plastics One, Roanoke, VA) were used to maintain the patency of the guide cannulae. During a 5-day recovery period, rats were extensively handled and repeatedly exposed to the manipulations associated with the microinjection procedure (BA et al., 2007). Prior to CIE exposure, all rats were microinjected with 0.5 μL/side aCSF to habituate them to the procedure. Correct placement of the guide cannulae was confirmed post-mortem by preparing coronal slices from fixed tissue.

### Chronic Intermittent Ethanol (CIE) Vapor Exposure

Pair-housed rats were exposed to chronic intermittent ethanol (CIE) vapor for 3 or 7 consecutive days, using standard procedures from our laboratory (Morales et al., 2018). Briefly, animals in their home cages were placed inside larger, custom-build Plexiglas chambers (Triad Plastics, Winston-Salem, NC). At the beginning of the light cycle (lights on at 9 PM), ethanol vapor was pumped into the chambers at a constant rate (16 L/min) and maintained at ~25 mg/L throughout the exposure for 12h/day. Air-exposed control animals were similarly housed, except they received room-air only. Animals were weighted daily, and tail blood samples were collected during the CIE exposure to monitor blood ethanol concentrations (BECs) and adjust ethanol vapor levels as necessary. Average BECs in the CIE animals were 241.29 ± 12.97 mg/dL as determined by a standard, commercially available alcohol dehydrogenase/NADH enzymatic assay (Carolina Liquid Chemistries, Greensboro, NC). These BECs are in the standard range for our laboratory (150-275 mg/dL). All behavioral assays and electrophysiology recordings were conducted 24h after the last ethanol (or air) exposure.

### Elevated Zero Maze

Twenty-four hours after the 7^th^ ethanol (or air) exposure, rats were microinjection with either artificial cerebral spinal fluid (aCSF; Tocris) or 300 μM clozapine-N-oxide (CNO; Tocris) and then tested on the elevated zero maze (EZM; Med Associates, Fairfax, VT) to assess anxiety-like behavior. The EZM is a circular maze consisting of two elevated closed or open sections, without the issue of an ambiguous center position found in the elevated plus maze. For microinjections, obturators were removed and 33 gauge injection cannulae were inserted into the guide cannulae such that their tips extended 1 mm below the ventral aspect of the guide cannulae. Drug solutions were bilaterally infused into the BLA over a 1 min period with 0.5 μL total volume delivered per side. Injectors were left in place for an additional minute before being removed. Animals were placed on the EZM 5 min after the microinjection (Poland et al., 2015). A Basler ace monochrome camera (Basler AG, Germany) along with EthoVision XT video tracking software (Noldus; Leesburg, VA) were used to track center point, tail base, and nose point throughout the 5 min test to assess anxiety-like behavior and general locomotion. Between animals, the apparatus was cleaned with warm water and mild soap, and then thoroughly dried.

### Electrophysiology

#### Slice Preparation

Animals were anesthetized with isoflurane and then decapitated using a guillotine. Brains were rapidly removed and incubated for 5 min in an ice-cold sucrose-modified artificial cerebral spinal fluid (aCSF) solution containing (in mM): 180 Sucrose, 30 NaCl, 4.5 KCl, 1 MgCl_2_·6H_2_O, 26 NaHCO_3_, 1.2 NaH_2_PO_4_, 10 D-glucose, 0.10 ketamine, equilibrated with 95% O_2_ and 5% CO_2_. Coronal slices (400 μM) containing the BLA were collected using a VT1200/S vibrating blade microtome (Leica, Buffalo Grove, IL) and incubated for ≥ 1h in room temperature (~25°C), oxygenated, standard aCSF solution containing (in mM): 126 NaCl, 3 KCl, 1.25 NaH_2_PO_4_, 2 MgSO_4_·7H_2_O, 26 NaHCO_3_, 10 D-glucose, and 2 CaCl_2_·2H_2_O prior to recordings. For low calcium recordings, an altered aCSF solution with 3 MgSO_4_ and 1 CaCl_2_ were used. All chemicals were obtained from Sigma-Aldrich (St. Louis, MO) or Tocris (Ellisville, Missouri).

#### Whole-Cell Patch-Clamp Recording

Methods for whole-cell voltage-clamp recordings were similar to those published previously from our laboratory (Morales et al., 2018). BLA slices were transferred to a submersion-type recording chamber that was continuously perfused at a rate of 2 mL/min with oxygenated, room temperature (~25°C) aCSF. Synaptic responses were recorded using electrodes filled with an intracellular solution containing (in mM): 145 CsOH, 10 EGTA, 5 NaCl, 1 MgCl2·6H2O, 10 HEPES, 4 Mg-ATP, 0.4 Na-GTP, 0.4 QX314, 1 CaCl_2_·2H_2_O, pH was adjusted to ~7.3 with gluconic acid, and osmolarity was adjusted to ~285 Osm/L with sucrose. Optogenetic (ChR2) responses were evoked using a 473 nm laser connected to a fiber optic cable (Thorlabs, Newton, NJ); the naked end of the cable was placed just above the stria terminalis on the medial side of the BLA. Five-millisecond laser pulses were used to activate channelrhodopsin found in the mPFC terminals. Sweeps were recorded every 30 s and light stimulation intensities were submaximal and normalized to elicit synaptic responses with amplitudes ~100 pA. Glutamatergic synaptic currents were recorded at a membrane holding potential of −70 mV and pharmacologically isolated using the GABA_A_ antagonist, picrotoxin (100 μM). GABAergic synaptic currents were recorded at a membrane holding potential of 0 mV and pharmacologically isolated with 20 μM 6,7-dinitroquinoxaline-2,3(1*H*,4*H*)-dione (DNQX, an AMPA/kainite receptor antagonist) and 50 μM DL-2-amino-5-phosphono-pentanoic acid (APV, an NMDA receptor antagonist). In some recordings, 1 μM tetrodotoxin (TTX; Tocris) and 20 mM 4-aminopyridine (4-AP; Tocris) were included for more stringent isolation of monosynaptic transmission (Cruikshank et al., 2010; Petreanu et al., 2009). Data were low-pass filtered at 2 kHz and acquired via an Axopatch 700B amplifier (Molecular Devices, Foster City, CA) and later analyzed using pClamp 10 software (Molecular Devises, Foster City, CA). Presumptive principal neurons were included based on their electrophysiological characteristics of high membrane capacitance (> 100 pF) and low access resistance (≤ 25 MΩ) (Washburn and Moises, 1992). Cells that did not meet these criteria or cells in which access resistance or capacitance changed ≥ 20% during the recording were excluded from analysis.

#### Paired-Pulse Ratio

Two 5-msec light stimuli of equal intensity were delivered to the stria terminalis at an inter-stimulus interval of 50-msec, with this short interval traditionally viewed as an indicator of presynaptic release probability (Fioravante et al., 2011). The paired-pulse ratio (PPR) was conservatively calculated using the evoked EPSC amplitudes as: ([Peak 2 amplitude – Peak 1 amplitude]/Peak 1 amplitude). The average paired-pulse ratio was determined for a 5-minute baseline and was compared to the average ratio of the final five minutes of drug application and washout.

### Experimental Design and Statistical Analysis

Primary statistical analyses were conducted using Prism 5 (GraphPad Software, La Jolla, CA). Data were analyzed using analysis of variance (ANOVA) or t-tests depending on the experimental design, with Bonferroni post doc tests used to determine the locus of effect as appropriate. A value of *p* < 0.05 was considered statistically significant. All data are presented as mean ± SEM throughout the text and figures.

## Results

### Both the PL/dmPFC and IL/vmPFC send robust projections to the BLA

Figure 1A shows the experimental timeline for the present study along with the approximate age of the animals at each phase. Figure 1B is a schematic demonstrating the approximate level of mPFC and BLA on a sagittal view of a rat brain: 1) mPFC, 2) BLA. Injection of AAV-ChR2-YFP targeting either the PL/dmPFC (Figure 1C) or IL/vmPFC (Figure 1D) results in robust YFP-expressing terminals in the BLA. Expression patterns between BLA-projecting PL/dmPFC and IL/vmPFC neurons were very similar across all levels of the BLA (data not shown). There were little to no YFP-expressing terminals in the lateral amygdala (LA) or the central nucleus of the amygdala (CeA), which is a typical innervation pattern for projections from the mPFC (McGarry and Carter, 2017; Reppucci and Petrovich, 2016). Figure 1E depicts the typical set-up for electrophysiology recordings. The fiber optic cable used to deliver 470 nm blue light was placed just outside of the BLA above the stria terminalis to activate the channelrhodopsin-expressing terminal entering the BLA from the mPFC. BLA principal neurons were patched with recording electrodes that were placed near the stimulating fiber where the YFP-expressing terminals were most dense.

**Figure 1.**
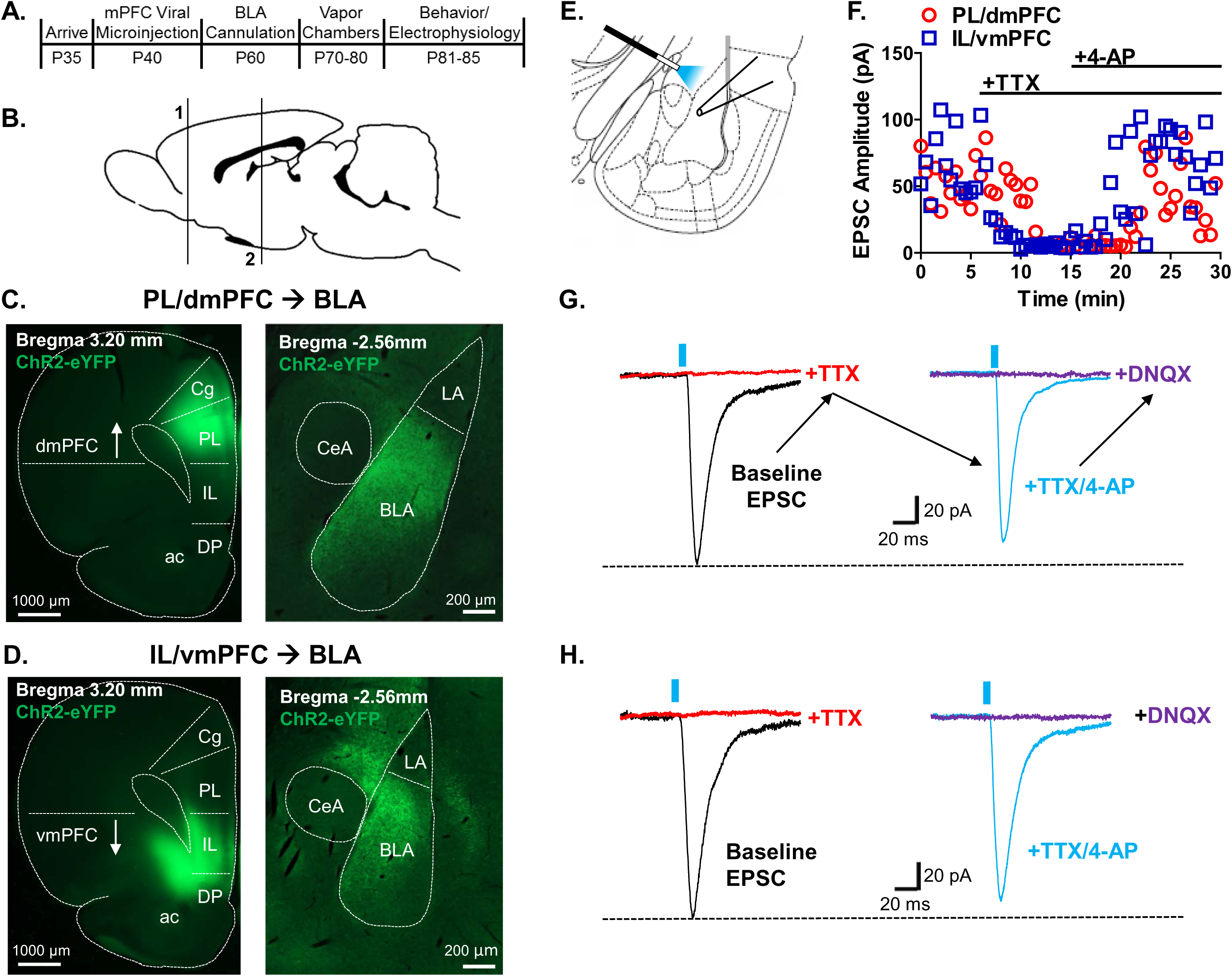
The dorsal and ventral subdivisions of the mPFC make monosynaptic glutamatergic synaptic connections with pyramidal neurons in the BLA. ***A***, Experimental timeline with major stages on the top and approximate age of the rat in postnatal days on the bottom. ***B***, Schematic of a sagittal view of a rat brain with the numbered lines demonstrating the relative locations of the brain regions examined including: 1) medial prefrontal cortex (mPFC) and 2) basolateral amygdala (BLA). ***C***, Representative fluorescent images of the prelimbic (PL) injection site (left) and the resulting terminals in the BLA (right) 4 weeks after injection of channelrhodopsin (ChR2). Note the arrow is depicting the anatomical location of what we refer to as the dorsal medial prefrontal cortex (dmPFC). ***D***, Representative fluorescent images of the infralimbic (IL) injection site (left) and the resulting terminals in the BLA (right) 4 weeks after injection of ChR2. The arrow is depicting the anatomical location of what we refer to as the ventral medial prefrontal cortex (vmPFC). ***E***, Schematic of our standard electrophysiology set-up with the recording pipette in the BLA and the optogenetic stimulating laser placed above the medial stria terminalis where mPFC inputs enter the BLA. ***F***, Glutamatergic excitatory postsynaptic current (EPSC) amplitude in picoamps (pA) as a function on time in minutes. The PL/dmPFC response amplitudes are denoted by circles and the IL/vmPFC response amplitudes are marked with squares. Bath application of tetrodotoxin (TTX; 1μM) abolishes light-evoked EPSCs, addition of 4-aminopyradine (4-AP; 20mM) rescues EPSCs. ***G***, Representative traces on light-evoked EPSCs recorded from PL/dmPFC – BLA and ***H***, IL/vmPFC – BLA synapses. The blue dashes represent optogenetic stimulation (5msec) with blue light. DNQX (20μM) abolished glutamate EPSC. Cg= cingulate cortex, PL= prelimbic cortex, IL= infralimbic cortex, DP= dorsal peduncular cortex, dmPFC= dorsal medial prefrontal cortex, vmPFC= ventral medial prefrontal cortex, ac= anterior commissure, CeA= central nucleus of the amygdala, LA= lateral amygdala, BLA= basolateral amygdala.

### Neurons from the PL/dmPFC and IL/vmPFC make monosynaptic glutamatergic synapses with principal neurons in the BLA

In order to test if these synaptic connections were monosynaptic, we recorded excitatory postsynaptic currents (EPSCs) in the presence of tetrodotoxin (TTX; sodium channel blocker) and 4-aminopyradine (4-AP; potassium channel blocker) (Figure 1F). After a stable baseline EPSC was collected, 1 μM TTX was applied until the response went away, approximately 5 minutes. With TTX still washing on, we added 1 mM 4-AP to the bath solution and the EPSCs returned after roughly 5 minutes. The amplitude of EPSCs collected in the presence of TTX and 4-AP were 89.54% ± 22.29 (PL/dmPFC) and 81.39% ± 11.64 (IL/vmPFC) of the baseline EPSC amplitudes, suggesting that a majority of the synaptic responses we recorded were monosynaptic, which others have also reported (Cho et al., 2013; Little and Carter, 2013). Once the EPSCs had returned in the presence of TTX and 4-AP, we washed on the AMPA receptor antagonist, DNQX (20 μM), to confirm that the responses were in fact glutamatergic. Application of DNQX abolished the monosynaptic EPSC. Figures 1G and 1H are traces from representative cells receiving inputs from the PL/dmPFC or IL/vmPFC, respectively.

### Withdrawal from 3 days of CIE does not alter glutamate release from mPFC terminals in the BLA

After confirming that we could reliably evoke and record monosynaptic glutamatergic EPSCs from PL/dmPFC and IL/vmPFC terminals, we wanted to examine the effects of chronic ethanol exposure and withdrawal on these circuits. Figure 2A is a timeline schematic depicting the 3- or 7-day CIE exposure that the ethanol-exposed animals received. Figure 2B shows that the glutamate release probabilities recorded from control air-exposed animals are significantly different (PL/dmPFC: −0.2615 ± 0.03735, N=21; IL/vmPFC: −0.6559 ± 0.03173, N=23) [Unpaired t-test (PL/dmPFC vs IL/vmPFC); t(42) = 8.090, p < 0.0001]. Next, we exposed animals to 3 days of CIE based on recently published data from our laboratory demonstrating that a single 3 day CIE exposure is sufficient to produce presynaptic changes (Morales et al., 2018). Surprisingly, withdrawal from 3 days of CIE exposure did not significantly alter the glutamate release probability from PL/dmPFC (AIR: −0.2960 ± 0.03712, N=11; CIE/WD: −0.3221 ± 0.05159, N=10) [Unpaired t-test (AIR vs CIE/WD); t(19) = 0.4170, p = 0.6814] or IL/vmPFC (AIR: −0.6344 ± 0.05184, N=10; CIE/WD: −0.5754 ± 0.08719, N=8) [Unpaired t-test (AIR vs CIE/WD); t(16) = 0.6094, p = 0.5508] terminal in the BLA (Figures 2C and 2D, respectively).

**Figure 2.**
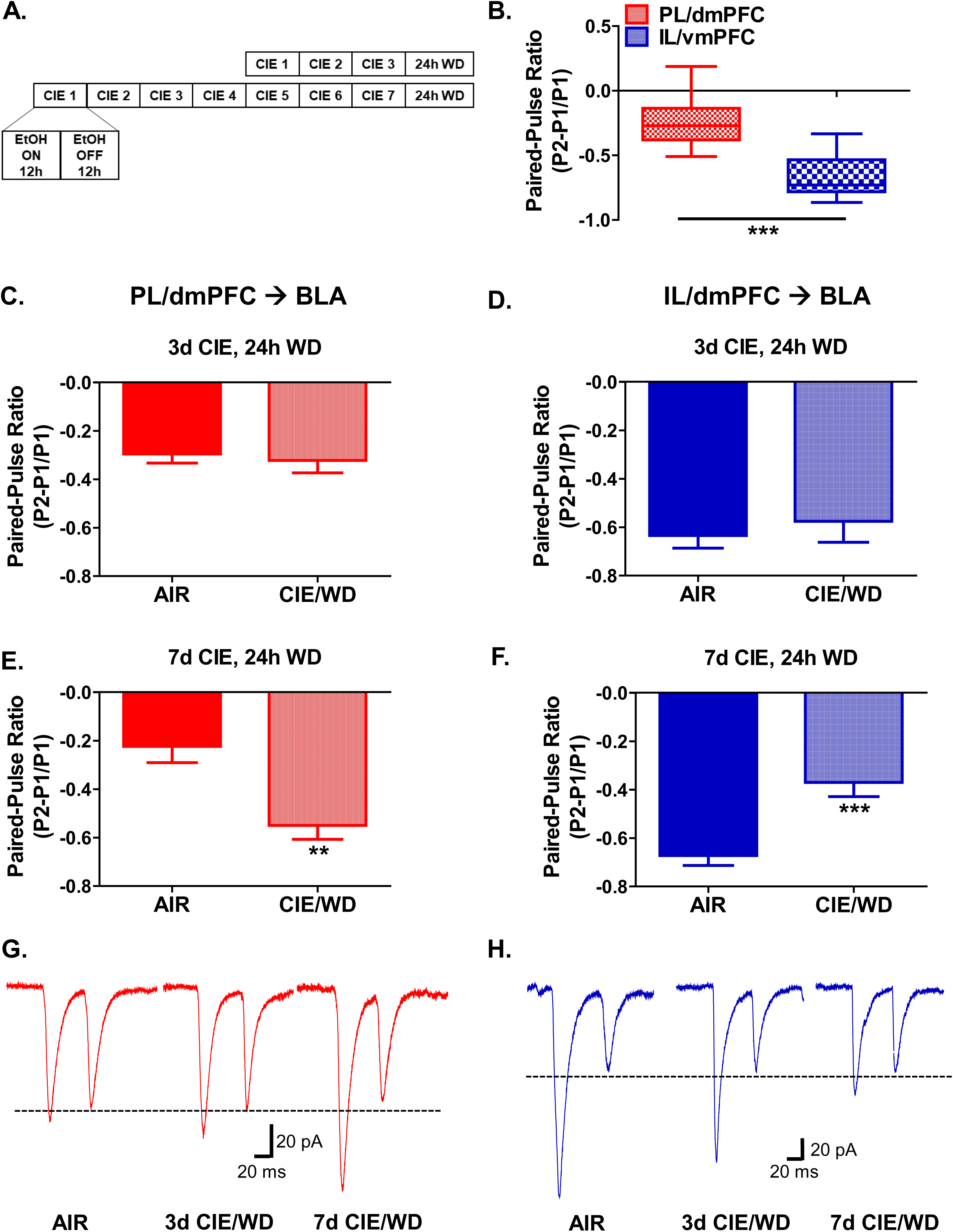
Withdrawal (WD) from 7, but not 3, days of CIE bidirectionally modulates glutamate release from PL/dmPFC and IL/vmPFC terminals in the BLA. ***A***, Timeline demonstrating the 3- and 7-day chronic intermittent ethanol (CIE) exposures. Each day, rats were exposed to 12- hours of ethanol (EtOH) and 12-hours of air. All electrophysiology recordings were conducted 24-hours after the last exposure. Control animals were only exposed to air. ***B***, Optogenetically-evoked glutamate paired-pulse ratios (PPR) recorded from BLA principal neurons receiving inputs from either the PL/dmPFC (N=21) or IL/vmPFC (N=23) were significantly different in controls. ***C***, No effect of 3-day CIE on PPR recorded from PL/dmPFC – BLA neurons in AIR (N=11) vs CIE/WD (N=10). ***D***, No effect of 3-day CIE on PPR recorded from IL/vmPFC – BLA neurons in AIR (N=10) vs CIE/WD (N=8). ***E***, Significant decrease in the PPR after 7-days of CIE in PL/dmPFC – BLA neurons, AIR (N=10) vs CIE/WD (N=8). ***F***, Significant increase in the PPR after 7-days of CIE in IL/vmPFC – BLA neurons, AIR (N=13) vs CIE/WD (N=15). ***G***, Representative traces of PPR recorded from PL/dmPFC – BLA or ***H***, IL/vmPFC – BLA neurons from AIR (left), 3-day CIE/WD (middle), and 7-day CIE/WD (right). **p<0.01; ***p<0.001; unpaired t-test.

### Withdrawal from 7 days of CIE bidirectionally modulates glutamate release from PL/dmPFC and IL/vmPFC terminals in the BLA

We then exposed animals to a longer, 7 day CIE exposure, which resulted in a significant increase in the paired-pulse ratios from PL/dmPFC terminals (AIR: −0.2237 ± 0.06725, N=10; CIE/WD: −0.5495 ± 0.05766, N=8) [Unpaired t-test (AIR vs CIE/WD); t(16) = 3.567, p = 0.0026] (Figure 2E) and a significant decrease in the paired-pulse ratio from IL/vmPFC terminals (AIR: − 0.6723 ± 0.04078, N=13: CIE/WD: −0.3704 ± 0.05866, N=15) [Unpaired t-test (AIR vs CIE/WD); t(26) = 4.100, p = 0.0004] (Figure 2F) in the BLA as compared to air-exposed control animals. As the paired-pulse ratio is inversely related to release probability, these data suggest that ethanol facilitates glutamate release from PL/dmPFC terminal and suppresses glutamate release from IL/vmPFC terminals. Figures 2G and 2H show normalized representative traces of the paired-pulse responses from air, 3-day CIE, and 7-day CIE animals recorded from BLA projecting PL/dmPFC and IL/vmPFC neurons, respectively. Interestingly, we noticed that PL/dmPFC and IL/vmPFC terminals had vastly different glutamate release probabilities at baseline (Figure 2B).

### Lowering calcium levels disrupts glutamate release from IL/vmPFC terminals but not PL/dmPFC terminals in the BLA

When examining the effects of chronic ethanol exposure on projections from the PL/dmPFC and IL/vmPFC, we found that the release probabilities in air-exposed control animals were significantly different (Figure 2B). Paired-pulse ratios are a measure of presynaptic release probability, and its long been known that calcium plays an important role in regulating neurotransmitter release (Neher and Sakaba, 2008). Therefore, we sought to examine the effects of lowering calcium levels on presynaptic glutamate release. Reducing calcium levels in the bath aCSF from 2 mM to 1mM had no effect on glutamate release probability from PL/dmPFC terminals in the BLA (Baseline: −0.3754 ± 0.02049; Low Calcium: −0.3798 ± 0.05923; Washout: −0.4820 ± 0.02923, N=14) (Figure 3A). However, the release probability from IL/vmPFC terminals was significantly decreased when calcium was lowered (Baseline: −0.6080 ± 0.04206; Low Calcium: −0.3515 ± 0.08208; Washout: −0.6551 ± 0.03278, N=15). A two-way repeated-measures ANOVA revealed a main effect of calcium levels [F(2,54) = 19.19, p < 0.0001], a main effect of brain region [F(1,54) = 4.74, p = 0.0416], and a significant interaction [F(2,54) = 8.55, p = 0.0006]. Bonferroni posttests (PL/dmPFC vs IL/vmPFC) further revealed a significant difference between the release probabilities from PL/dmPFC and IL/vmPFC terminals at baseline [t = 3.318, p < 0.01] and during washout [t = 2.470, p < 0.05], when the calcium levels were normal (Figure 3A). Interestingly, lowering calcium levels decrease IL/vmPFC release probability to PL/dmPFC levels to the point where they were not longer significantly different [t = 0.4048, p > 0.05]. Figure 3B shows normalized release probabilities so that we could quantify the magnitude of effect of altering calcium levels. Glutamate release from IL/vmPFC terminals was reduced by 42.19% ± 13.50, with no significant effect on PL/dmPFC release (101.2% ± 15.78). These data were analyzed using a two-way repeated-measure ANOVA, which revealed a main effect of calcium level [F(1,27) = 7.357, p = 0.0115] and a significant interaction [F(1,27) = 8.227, p = 0.0079]. Planned Bonferroni posttests (Baseline vs Low Calcium) further revealed a significant difference in the effect of calcium levels on brain region such that PL/dmPFC was unchanged [t = 0.1084, p > 0.05] but IL/vmPFC was significantly different [t = 4.016, p < 0.001]. Representative traces of paired-pulse responses recording during baseline (2 mM Ca^2+^), low calcium (1 mM Ca^2+^), and washout (2 mM Ca^2+^) from PL/dmPFC and IL/vmPFC terminals in the BLA are shown in Figures 3C and 3D, respectively.

**Figure 3.**
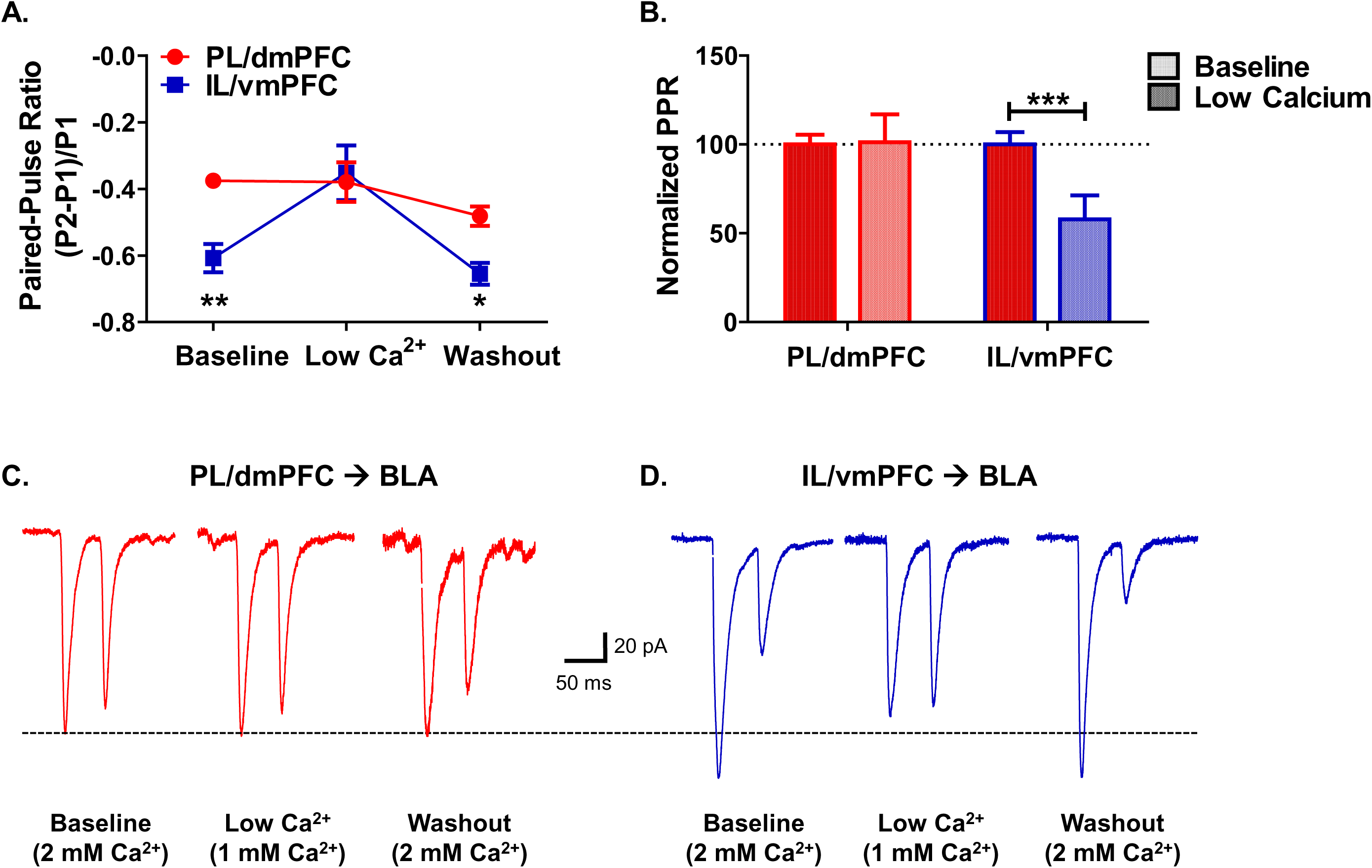
Lowering calcium levels disrupts glutamate release from IL/vmPFC terminals but not PL/dmPFC terminals in the BLA. ***A***, Paired-pulse ratios recorded from PL/dmPFC or IL/vmPFC terminals in the BLA during baseline (2mM Ca^2+^), low calcium (1mM Ca^2+^), and washout (2mM Ca^2+^) in the bath solution in slices from air-exposed control aniamls. Significant differences in the PPR between PL/dmPFC (N=14) and IL/vmPFC (N=15) at baseline and washout but not in the presence of low calcium. ***B***, Normalized PPR with the low calcium PPRs calculated as a percentage of baseline PPR allowing for direct comparison of magnitude of effect of lower calcium levels on glutamate release probability. Significant differences between baseline and low calcium for IL/vmPFC synapses but not PL/dmPFC. ***C***, Representative traces of light-evoked paired-pulse responses with a 50msec interstimulus interval from PL/dmPFC – BLA projections and ***D***, IL/vmPFC – BLA projections with baseline (left), low calcium (middle), and washout (right). *p<0.05; **p<0.01; ***p<0.001; Two-way repeated measures ANOVA.

### Bath application of CNO decreases glutamate release from Gi-DREADD expressing PL/dmPFC terminals in the BLA

Since we know that increased glutamate release in the BLA can lead to increases in anxiety-like behavior (McCool et al., 2010), we wanted to use DREADD technology to selectively decrease glutamate release from PL/dmPFC terminals in the BLA. Approximately 6 weeks after co-injecting channelrhodopsin (ChR2) and Gi-DREADD containing viruses into the PL/dmPFC, we examined the effects of 10 µM clozapine-N-oxide (CNO) bath application on paired-pulse ratios recorded from BLA principal neurons. We found that CNO significantly decreased the glutamate paired-pulse ratio in both AIR (Baseline: −0.2604 ± 0.05714; CNO: −0.04875 ± 0.08127, N=6) [Paired t-test (Baseline vs CNO); t(5) = 3.70, p = 0.0139] (Figure 4A) and CIE/WD animals (Baseline: −0.6248 ± 0.02394; CNO: −0.3777 ± 0.03984, N=6) [Paired t-test (Baseline vs CNO); t(5) = 6.35, p = 0.0014] (Figure 4B). Importantly, CNO had no effect on glutamate paired-pulse ratios in recordings from animals (AIR and CIE/WD combined) lacking the Gi-DREADD (“ChR2 only”) (Baseline: −0.5411 ± 0.04300; CNO: -.05598 ± 0.04969, N=11) [Paired t-test (Baseline vs CNO); t(10) = 0.98, p = 0.3456] (Figure 4C). Representative traces of paired-pulse ratios recorded at baseline and in the presence of CNO from AIR and CIE/WD animals are shown in Figures 4D and 4E, respectively. Additionally, the percent inhibition of peak 1 was similar in both AIR (68.32 ± 14.09, N=6) and CIE/WD (59.41 ± 12.77, N=6) animals [Unpaired t-test (AIR vs CIE/WD); t(10) = 0.4684, p = 0.6496] (Figure 4F).

**Figure 4.**
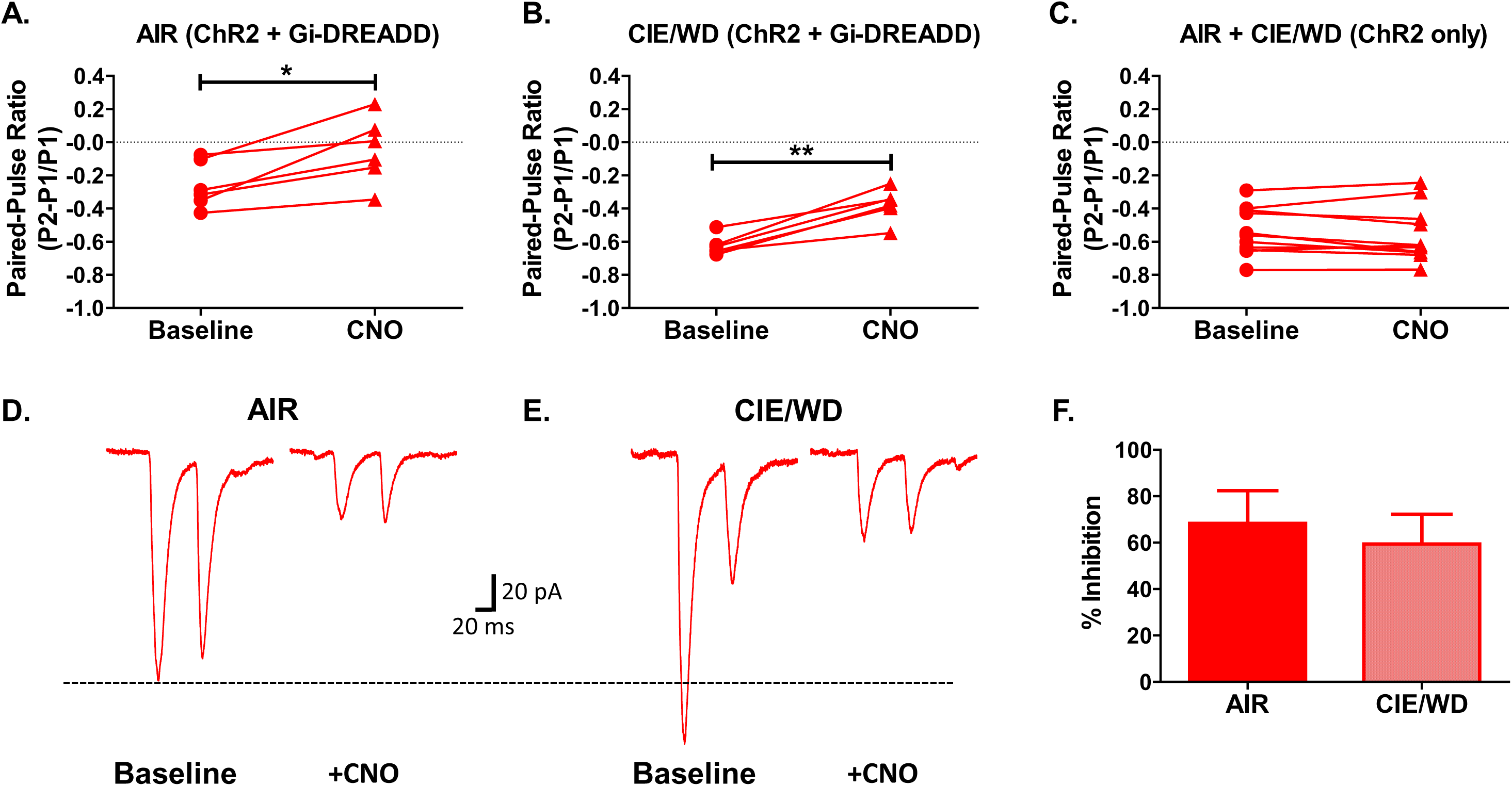
Bath application of CNO decreases glutamate release from Gi-DREADD expressing PL/dmPFC terminals in the BLA. ***A***, Light-evoked paired-pulse ratios recorded from PL/dmPFC – BLA terminals expressing ChR2 and Gi-coupled DREADDs at baseline and during the bath application of CNO (10μM) in air-exposed control animals (N=6). CNO significantly increased the PPR in all the cells recorded. ***B***, PPR recorded from CIE/WD animals (N=6) expressing ChR2 and Gi-DREADDs with and without CNO. Bath application of CNO significantly increased the PPR in all cells recorded. ***C***, PPRs recorded from AIR and CIE/WD animals (N=11) with ChR2 only. Bath application of CNO did not significantly change glutamate release probability from terminals without the DREADD construct. ***D***, Representative traces of paired-pulse responses recorded from AIR or ***E***, CIE/WD animals during baseline (left) and in the presence on CNO (right). ***F***, No difference in the percent inhibition of PPR peak 1 amplitude between AIR and CIE/WD recordings. *p<0.05; **p<0.01; Paired t-test.

### Chemogenetic inhibition of glutamate release from PL/dmPFC terminals in the BLA decreases anxiety-like behavior during withdrawal

After demonstrating that activation of Gi-coupled DREADDs with CNO decreases glutamate release from PL/dmPFC terminals in the BLA, we sought to examine the behavioral consequences of silencing this PL/dmPFC – BLA circuit during ethanol withdrawal. Figure 5A contains images taken using a fluorescent microscope depicting mCherry expression in neurons at the injection site, in the PL/dmPFC. Figure 5B is a schematic demonstrating the areas of the BLA that were targeted with the chronic cannulation implantation for CNO microinjection. Black dots represent correct cannula placements, grey dots represent incorrect cannula placements for which animals were excluded from analysis (N=4). Our target level of BLA was AP −2.80 mm from bregma, and all of the cannulations fell between −2.56 mm and −3.14 mm from bregma which was revealed through post-hoc analysis of sliced rat brains. We replicated and extended previous findings from our lab (Morales et al., 2018) by demonstrating that withdrawal from 7 days of CIE produces an increase in anxiety-like behavior measured by percentage of time spent in the open areas on the elevated zero maze (AIR ACSF: 16.15% ± 3.898, N=8; CIE/WD ACSF: 3.962% ± 1.636, N=8) (EZM; Figure 5C). Percentage of time was calculated by dividing the time (in seconds) spent collectively in both of the open areas by the entire time of the test, which was 300 seconds. Importantly, in rats with the Gi-coupled inhibitory DREADDs, microinjection of CNO (300 µM) into the BLA significantly increased the time that ethanol exposed animals spent in the open areas of the EZM, suggesting a decrease in anxiety-like behavior (CIE/WD CNO: 16.98% ± 4.368, N=7) (Figure 5C). These data were analyzed using a two-way ANOVA, which revealed a significant interaction (exposure X treatment) [F(1,27) = 9.185, p = 0.0053]. Planned post-hoc analyses using Bonferroni’s posttest demonstrated a significant withdrawal effect (AIR vs CIE) only for animals in the ACSF treatment group [t=2.564, p < 0.05], but not for animals treated with CNO [t=1.737, p > 0.05]. Further comparisons of treatment effect (ACSF vs CNO) showed a significant effect of CNO in the CIE/WD animals [t=2.647, p < 0.05], but no significant effect of CNO in AIR-exposed control animals (AIR CNO: 8.440% ± 3.368, N=8) [t=1.621, p > 0.05] (Figure 5C). We used total distance moved as a measurement of locomotor activity on the EZM (Figure 5D). A two-way ANOVA revealed a main effect of exposure group (AIR vs CIE/WD) [F(1,27) = 7.194, p = 0.0123] and a significant interaction (exposure X treatment) [F(1,27) = 7.334, p = 0.0116]. Total distance, in millimeters, is as follows: AIR ACSF: 2,629 ± 340.5, N=8; AIR CNO: 1,731 ± 183.0, N=8; CIE/WD ACSF: 1,402 ± 185.8, N=8; CIE/WD CNO: 1,737 ± 110.4, N=7. Figure 5E contains EZM heat plots based on group means, with white/lighter color representing less time and red representing more time.

**Figure 5.**
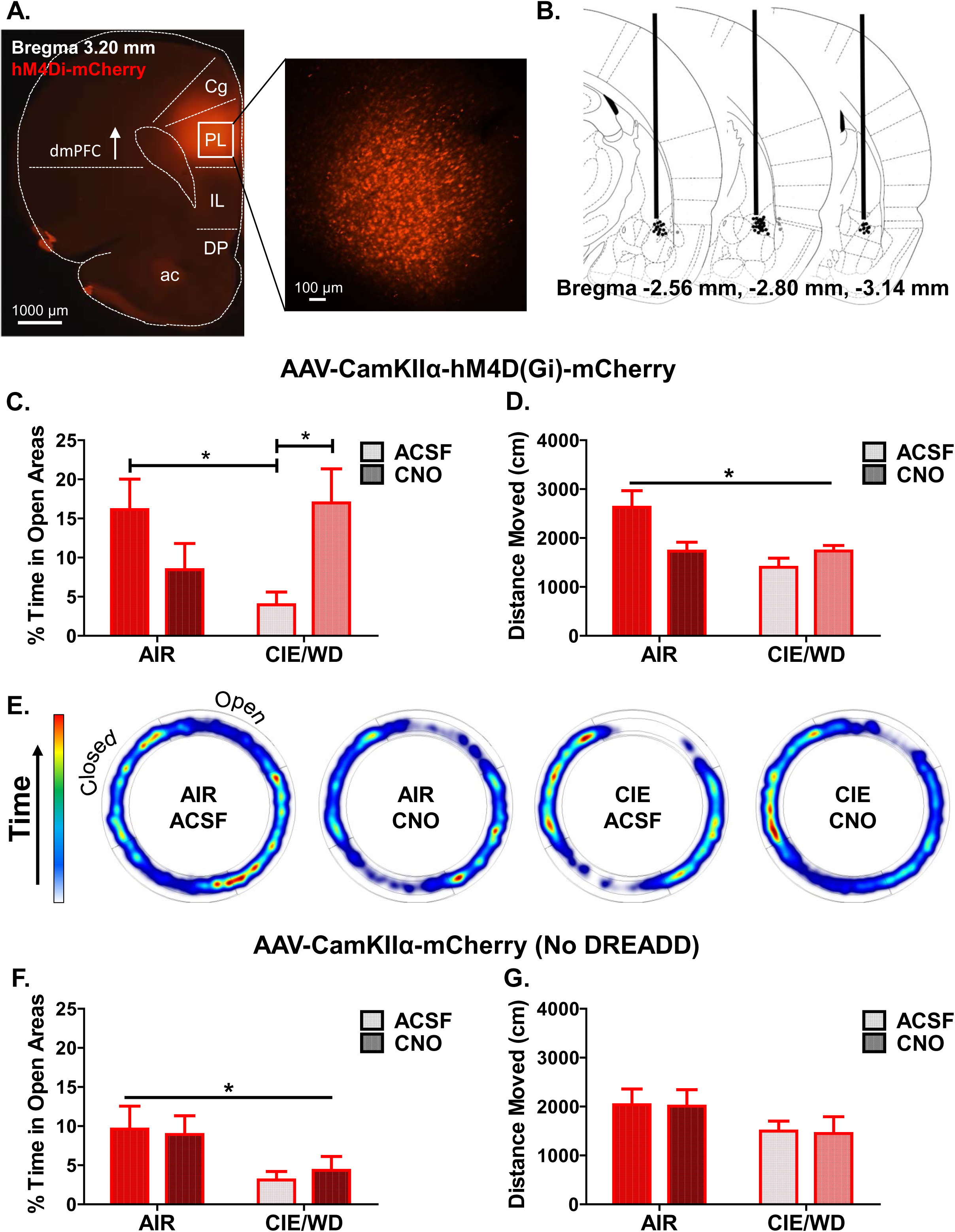
Selectively inhibiting glutamate release from PL/dmPFC terminals in the BLA attenuates withdrawal-induced anxiety-like behavior. ***A***, Representative fluorescent images of the PL-targeted injection site for the Gi-coupled DREADDS (mCherry). ***B***, Schematic depicting level of the BLA targeted for cannula placement relative to bregma. Black dots represent approximate location of cannula tip that fell within the region of interested and were included in the analysis. Grey dots represent cannula tip placements that fell outside of the BLA and were excluded from analysis. ***C***, Percent time spent in the open areas of the elevated zero maze for AIR (N=16) and CIE/WD (N=15) animals expressing Gi-DREADDs that received microinjections of either ACSF (N=8/group) or CNO (N=7-8/group; 300μM). Significant difference between AIR ACSF vs CIE/WD ACSF and between CIE/WD ACSF vs CIE/WD CNO. No significant differences between AIR ACSF vs AIR CNO nor AIR ACSF and CIE/WD CNO. ***D***, Main effect of exposure condition (AIR vs CIE/WD) for total distance moved in centimeters (cm) on the EZM. ***E***, Representative heat plots of amount of time spent in open vs closed areas of the EZM depicting group data. ***F***, Main effect of exposure condition (AIR vs CIE/WD) on percentage of time spent in the open areas of the EZM in animals without the DREADD construct (N=4/group). ***G***, No differences observed between any of the conditions for total distances moved. *p<0.05, Two-way ANOVA. Cg= cingulate cortex, PL=prelimbic cortex, IL= infralimbic cortex, DP= dorsal peduncular cortex; ac= anterior commissure, dmPFC= dorsal medial prefrontal cortex.

### CNO has no effect in rats injected with the control virus lacking the Gi-coupled DREADD

We generated another group of animals that were injected with a mCherry control virus lacking the Gi-DREADD construct. We used these animals to examine the effects of microinjecting CNO into the BLA to see if CNO alone would produce any behavioral alterations (Figure 5F and 5G). A two-way ANOVA of percent time spent in open area revealed a main effect of exposure (AIR vs CIE/WD) [F(1,12) = 6.627, p = 0.0244], with no main effect of treatment (ACSF vs CNO) [F(1,12) = 0.01565, p = 0.9025] and no interaction (exposure X treatment) [F(1,12) = 0.1968, p = 0.6652] (Figure 5F). Percent time spent in open areas for animals with control virus is as follows: AIR ACSF: 29.45% ± 8.973, N=4; AIR CNO: 27.35% ± 7.321, N=4; CIE/WD ACSF: 9.55% ± 3.274, N=4; CIE/WD CNO: 13.30% ± 5.395, N=4. Again, we used total distance traveled, in millimeters, as a measurement of locomotor activity on the EZM. A two-way ANOVA of total distance traveled revealed no main effects or interactions: exposure condition [F(1,12) = 3.22, p = 0.0979], treatment group [F(1,12) = 0.01578, p = 0.9021], interaction [F(1,12) = 0.001529, p = 0.9695] (Figure 5G). Total distance, in millimeters, is as follows: AIR ACSF: 2,036 ± 321.2, N=4; AIR CNO: 2,010 ± 334.8, N=4; CIE//WD ACSF: 1,498 ± 205.1, N=4; CIE/WD CNO: 1,447 ± 345.0, N=4.

### Withdrawal from CIE increases the E/I ratio in the PL/dmPFC – BLA pathway

To gain a better understanding of the PL/dmPFC to BLA pathway and the alterations produced by chronic ethanol exposure, we examined the balance between excitatory and inhibitory transmission evoked by PL/dmPFC terminals in the BLA. In the presence of the NMDA receptor antagonist, APV (50 µM), we recorded optically evoked glutamatergic EPSCs at −70 mV followed by GABAergic IPSCs at 0 mV. Since we have already shown that withdrawal from chronic ethanol increase glutamate transmission at these synapses (Figure 2E), we adjusted the laser intensity to produce equivalent EPSCs and then recorded the resulting IPSC from that same stimulation intensity as a way to normalize stimulation intensities between cells. We found that withdrawal from 7 days of CIE significantly increases the EPSC/IPSC ratio (AIR: 0.6326 ± 0.08788, N=6; CIE/WD: 1.536 ± 0.266, N=9) [Unpaired t-test (AIR vs CIE/WD); t(13) = 2.677, p = 0.019] (Figure 6A). Further examination of the peak amplitudes of the EPSCs (AIR: 154.2 ± 32.27, N=6; CIE/WD: 123.8 ± 12.54, N=10) and IPSCs (AIR: 278.1 ± 83.38, N=6; CIE/WD: 130.4 ± 29.97, N=10) revealed that this increase in the E/I ratio was driven by a decrease in the average peak amplitude of IPSCs recorded from CIE/WD animals (Figure 6B). A two-way ANOVA revealed a main effect of exposure (AIR vs CIE) [F(1,26) = 5.994, p = 0.0214]. There was no main effect of response type (EPSC vs IPSC) [F(1,26) = 1.743, p = 0.1982] and no significant interaction (exposure X response) [F(1,26) = 3.28, p = 0.0817]. Bonferroni’s posttest (AIR vs CIE/WD) indicated a significant difference in the IPSC peak amplitude [t = 3.012, p < 0.05] with no significant different in EPSC peak amplitude [t = 0.4505, p > 0.05] (Figure 6B). Interestingly, the latency of response onset, measured in milliseconds post-stimulation, was significantly longer for evoked IPSCs (AIR: 62.52 ± 2.239, N=6; CIE/WD: 64.97 ± 1.632, N=10) than EPSCs (AIR: 50.52 ± 0.7593, N=6; CIE/WD: 50.01 ± 0.2947, N=10), regardless of exposure condition (Figure 6C). Importantly, EPSCs were sensitive to the glutamate antagonist, DNQX (20 µM) but not the GABA_A_ antagonist, picrotoxin (100 µM), or the combination to TTX (1 µM) and 4-AP (1mM). IPSCs were abolished by application of picrotoxin, DNQX, and TTX +4- AP (data not shown).

**Figure 6.**
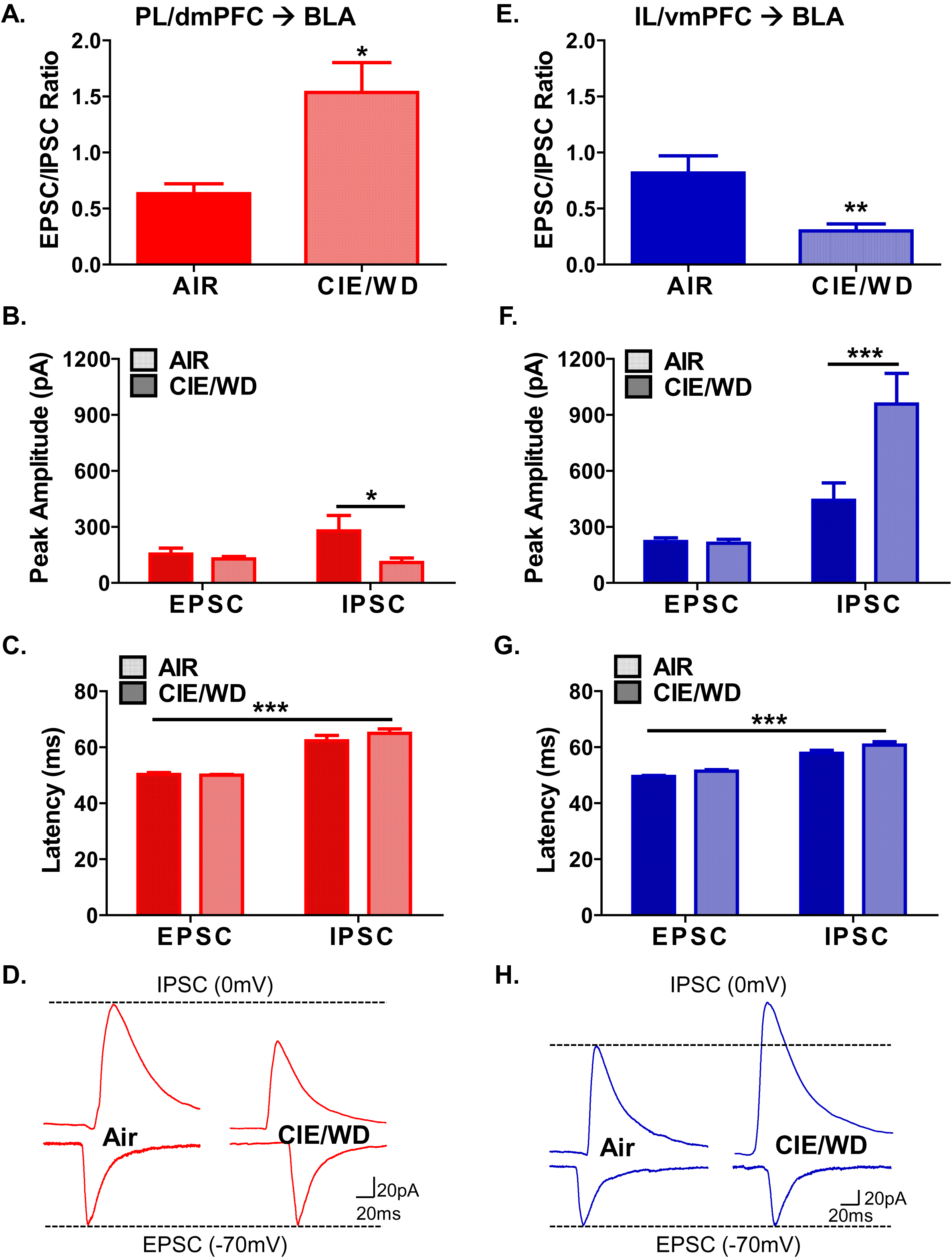
Withdrawal from CIE produces opposing alterations to the balance of excitation and inhibition in the PL/dmPFC and IL/vmPFC – BLA pathway. ***A***, Light-evoked EPSC/IPSC (E/I) ratio recorded from PL/dmPFC – BLA terminals. Significant increase in the E/I ratio in CIE/WD (N=9) vs AIR (N=6). ***B***, Peak amplitudes in picoamps (pA) of glutamatergic EPSCs and GABAergic IPSCs recorded from PL/dmPFC – BLA synapses. Significant reduction in the IPSC amplitude in CIE/WD animals. ***C***, Response onset latency for EPSCs and IPSCs recorded from PL/dmPFC – BLA inputs. ISPCs had significantly longer latencies than EPSCs regardless of exposure condition. ***D***, Representative traces recorded from PL/dmPFC – BLA terminals. IPSC recorded at 0mV and EPSCs recorded at −70mV. ***E***, Light-evoked E/I ratios recorded from IL/vmPFC – BLA terminals. Significant decrease in the E/I ratio in CIE/WD (N=13) vs AIR (N=13). ***F***, Peak amplitudes of EPSCs and IPSCs recorded from IL/vmPFC – BLA synapses. Significant increase in the IPSC amplitude in CIE/WD animals. ***G***, Response onset latency for EPSCs and IPSCs recorded from IL/vmPFC – BLA inputs. ISPCs had significantly longer latencies than EPSCs regardless of exposure condition. ***H***, Representative traces recorded from IL/vmPFC – BLA terminals. *p<0.05; **p<0.01; ***p<0.001; Unpaired t-test or Two-way ANOVA.

### Withdrawal from CIE decrease the E/I ratio in the IL/vmPFC – BLA pathway

We then preformed the same experiment to look at the alteration in glutamatergic and GABAergic transmission evoked in BLA principal neurons by optically activating IL/vmPFC terminals. In contrast to what we found in the PL/dmPFC to BLA pathway, withdrawal from 7 days of CIE significantly decreased the EPSC/IPSC ratio in the IL/vmPFC to BLA pathway (Figure 6E). An unpaired t-test was used to compare the E/I ratios from AIR (0.8182 ± 0.1519, N=13) and CIE/WD (0.3013 ± 0.06194) animals [t(24) = 3.152, p = 0.0043]. Examination of the peak amplitudes (pA) demonstrated that this decrease in the E/I ratio was driven by a significant increase in the average IPSC amplitude recorded from CIE/WD animals (Figure 6F). These data were analyzed using a two-way ANOVA, which revealed a main effect of exposure (AIR vs CIE/WD) [F(1,48) = 7.030, p = 0.0108], a main effect of response (EPSC vs IPSC) [F(1,48) = 25.63, p < 0.0001], and a significant interaction (exposure X response) [F(1,48) = 7.568, p 0.0084] (Figure 6F). Bonferroni’s posttest further revealed that the IPSC peak amplitude was significantly different in AIR vs CIE/WD animals [t = 3.820, p < 0.001]. Importantly, the EPSC amplitudes were not significantly different [t = 0.0705, p > 0.05], indicating our standardization was successful. Interestingly, Bonferroni’s posttest also showed that in CIE/WD [t = 5.525, p < 0.001], but not AIR [t = 1.635, p > 0.05] animals, the EPSC (AIR: 221.7 ± 20.10, N=13; CIE/WD: 212.2 ± 20.89, N=13) and IPSC (AIR: 442.5 ± 93.15, N=13; CIE/WD: 958.6 ± 164.3, N=13) peak amplitudes were significantly different (Figure 6F). Similar to the PL/vmPFC pathway, the latencies (ms) of IPSCs (AIR: 57.82 ± 1.059, N=13; CIE/WD: 60.76 ± 1.202, N=13) were significantly longer than those of EPSCs (AIR: 49.54 ± 0.3721, N=13; CIE/WD: 51.40 ± 0.5569, N=13) recorded from stimulating the IL/vmPFC pathway (Figure 6G). These data were analyzed using a two-way ANOVA, which revealed a main effect of response (EPSC vs IPSC) [F(1,48) = 103.2, p < 0.0001], a main effect of exposure (AIR vs CIE/WD) [F(1,48) = 7.646, p = 0.0081], with no interaction (response X exposure) [F(1,48) = 0.3777, p = 0.5417]. Similarly to PL/dmPFC response, EPSCs evoked from IL/vmPFC terminals were sensitive to DNQX but remained intact in the presence of picrotoxin and TTX + 4-AP. Conversely, IPSCs were attenuated with the application of picrotoxin, DNQX, or TTX and 4-AP (data not shown).

## Discussion

The present study sought to examine the effects of chronic ethanol exposure and withdrawal on projections from two subdivisions of the medial prefrontal cortex to the basolateral amygdala. We used a combination of optogenetics and chemogenetics to document the neurophysiological and behavioral alterations of PL/dmPFC and IL/vmPFC terminals in the BLA. The major findings of the study are that withdrawal from chronic ethanol exposure increases optogenetically stimulated glutamate release from PL/dmPFC terminals but decreases the glutamate release from IL/vmPFC terminals in the BLA. Additionally, we demonstrate that chemogenetic inhibition of PL/dmPFC terminals in the BLA rescues withdrawal-induced anxiety-like behavior. Together, these data provide the first characterization of how chronic ethanol exposure and withdrawal changes mPFC – BLA circuitry and suggests a novel role for this circuit in regulating ethanol withdrawal-associated anxiety.

Through precise stereotaxic microinjection, we targeted the prelimbic (PL) and infralimbic (IL) subdivisions of the medical prefrontal cortex (mPFC). To prevent unintended viral spread between the PL and IL regions, we placed the tip of our injectors in the top third of the PL and the bottom third of the IL. This allowed for maximal separation of PL and IL but caused spread of the virus into the anterior cingulate (Cg) and dorsal peduncular (DP) cortices in some cases. Recently, the anterior cingulate cortex and its input to the basolateral amygdala has been implicated in mediating innate freezing response to predator odors (Jhang et al., 2018), which is similar to the role the PL plays in learned fears (Corcoran and Quirk, 2007). Although the projection from the dorsal peduncular cortex to the BLA has not been directly characterized, the DP is thought to play a role similar to the IL and is often included when examining conditioned responses such as fear and drug seeking (Peters et al., 2009). Hence, we refer broadly to the PL as the dorsal component (dmPFC) and IL as the ventral component (vmPFC). These are common subdivisions based on cytoarchitecture, functional criteria, and connectivity with other brain regions (Heidbreder and Groenewegen, 2003). Notably, homologies between the human, primate, and rodent prefrontal cortices exist, which allows for cross-species comparisons (Laubach et al., 2018). The pattern of terminal expression in the BLA was similar between PL/dmPFC and IL/vmPFC injected animals such that only cells in the basolateral nucleus were innervated and this was consistent along the rostral – caudal axis of the. In particular, the lateral nucleus and central nucleus of the amygdala were devoid of innervation BLA. These findings are consistent with previously reported terminal formation and holds true throughout development (Arruda-Carvalho et al., 2017).

The use of TTX and 4-AP to demonstrate monosynaptic circuits is imperative when using optogenetics and electrophysiology. Without these pharmacological agents, there is no way to distinguish if a light evoked response is coming from virally transfected fibers that make synapses directly onto the cell you are recording from or if the responses are from other cells in the local network. In the present study, application of TTX, which inhibits action potential firing, completely blocked optogenetically-evoked EPSCs recorded in the BLA that originated in both the PL and IL. The addition of 4-AP and subsequent blockade of voltage-gated potassium channels induced additional depolarization at the terminals so that ChR2-mediated depolarization was then enough to trigger glutamate release (Figure 1). Our results are consistent with the monosynaptic nature of the mPFC – BLA projections that has demonstrated previously by others (Arruda-Carvalho and Clem, 2014; Cho et al., 2013; Kiritoshi and Neugebauer, 2018; Little and Carter, 2013; McGarry and Carter, 2017). Importantly, we compared paired-pulse ratios recorded in the presence of TTX and 4-AP as well as under standard conditions and we did not observe a significant difference, suggesting that our recordings did not contain significant polysynaptic contributions to our glutamatergic recordings. In contrast, 4-AP failed to rescue TTX-blocked IPSCs suggesting the GABAergic responses recorded were disynpatic in nature (Figure 6).

A main focus of our laboratory is on understanding the synaptic changes that occur in response to chronic ethanol exposure and withdrawal. Previous studies from our laboratory have characterized the presynaptic and postsynaptic adaptions that occur following withdrawal from chronic ethanol exposure. For example, electrophysiology recordings using electrical stimulation of medial stria terminalis inputs have shown that withdrawal from a 10 day CIE exposure increase the glutamate release probability at these synapses (Christian et al., 2013) and that these presynaptic changes are present after just 3 days of CIE (Morales et al., 2018). Accordingly, we wanted to investigate if these findings held true for specific circuits. Projections from the mPFC to the BLA arrive via the medial stria terminalis (Sesack et al., 1989) so we hypothesized that these synapses would also undergo presynaptic alterations following withdrawal from chronic ethanol exposure. These data presented here demonstrate that mPFC synapses undergo presynaptic changes, but not necessarily in the manor we expected. For instance, withdrawal from 3 days of CIE did not alter the glutamate release probability at PL/dmPFC nor IL/vmPFC synapses in the BLA (Figure 2). Interestingly, withdrawal from 7 days of CIE significantly increased the release probability at PL/dmPFC synapses but decreased the release probability at IL/vmPFC synapses in the BLA relative to air-exposed control animals. We believe a plausible explanation for a lack of effect after 3 days of CIE in these circuits, which contrasts with the recently published study from our laboratory, is related to the age of the animals used in the different studies. Animals in our previous report were exposure to ethanol during adolescence (postnatal day 39-46), whereas animals used in the present study were adults at the time of ethanol exposure (P70+). Others have also found that adolescents are more vulnerable to the effects of ethanol and that these alterations can persist into adulthood (reviewed in (Crews et al., 2016)). Therefore, a plausible explanation for a lack of ethanol effect after the 3-day CIE exposure is that adults more resilient and therefore require a longer exposure to induce the same types of synaptic alterations.

We find that withdrawal from 7 days of CIE produces opposite effects on PL/dmPFC and IL/vmPFC projections to the BLA. Indeed, others have also reported differential regulation of these two pathways. For instance, fear learning leads to strengthening of PL but not IL excitatory synapses in BLA principal neurons (Arruda-Carvalho and Clem, 2014). Additionally, inhibition-excitation ratios in BLA neurons were increased in an arthritis pain model in the IL pathway but not in the PL pathway (Kiritoshi and Neugebauer, 2018). Taken together, these findings highlight that these two mPFC – BLA circuits likely serve independent functions and can be differentially altered under pathological conditions. As mentioned previously, the PL/dmPFC and IL/vmPFC play distinct roles in both fear and drug-seeking behaviors such that the PL/dmPFC drives the expression of fear and drug seeking, whereas the IL/vmPFC suppresses these behaviors after extinction (reviewed in (Peters et al., 2009)). Moreover, there’s a large body of evidence that suggests the mechanisms underlying fear and anxiety processes are mediated by overlapping neuronal circuits in both rats and humans (review in (Davis et al., 2010; Tovote et al., 2015). Based on the literature presented here, a possible functional consequence of our findings demonstrating changes in glutamate signaling in the BLA is that the increase in glutamate release from PL/dmPFC terminals during alcohol withdrawal acts to promote fear, anxiety, and drug seeking whereas the observed decrease in glutamate release from IL/vmPFC terminals decrease the ability of these projections to suppress these behaviors, ultimately leading to the pathological state observed during abstinence.

The BLA has been implicated in mediating a wide range of behaviors (reviewed in (Janak and Tye, 2015)) and increases in glutamate signaling within the BLA specifically is associated with increases in emotional behaviors such as fear and anxiety (reviewed in (McCool et al., 2010)). On the other hand, decreases in glutamate release has been linked to an anxiolytic state, and our lab has previously demonstrated that microinjection of the glutamate receptor antagonist, DNQX, into the BLA attenuates anxiety-like behavior expressed during withdrawal (Läck et al., 2007). In line with this, our current findings show similar results such that microinjection of CNO into the BLA diminished the increased anxiety-like behavior seen during withdrawal from 7 days of CIE (Figure 5). This effect was mediated through activation of Gi-coupled DREADDs expressed in PL/dmPFC terminals in the BLA, which we’ve shown effectively suppresses glutamate release at these synapses in both AIR and CIE/WD animals (Figure 4). Our EZM data replicated previous findings of ours that withdrawal from 7 days of CIE exposure increase anxiety like behavior and extended the current literature by directly demonstrating that PL/dmPFC projections to the BLA play an important role in regulating anxiety-like behavior. This is of particular importance because increased negative affect during withdrawal has been reported as a common underlying cause of relapse in human alcoholics (Willinger et al., 2002). Although projections from the BLA to the mPFC have been shown to bidirectionally modulate anxiety-related behaviors (Felix-Ortiz et al., 2015), the ability of mPFC projections to the BLA to do the same has not been explored prior to this study. We observed increased glutamate release from PL/dmPFC terminals in the BLA during withdrawal and we found that inhibiting this glutamate release in vivo using the Gi-coupled DREADD attenuated the increased-anxiety-like behavior. Therefore, we also hypothesized that increasing glutamate release from IL/vmPFC terminals in the BLA would rescue the increased anxiety-like behavior since we see a reduction during alcohol withdrawal. Also, electrical stimulation of IL has been shown to increase the time spent and entries in the open arms of an elevated-plus maze suggesting that the IL plays a key role in exerting active action to overcome anxiety-like behavior (Shimizu et al., 2018), which parallels the role this region plays in extinction learning. We tried to test this question by expression Gq-coupled DREADDs in the IL/vmPFC, however, all our animals that we injected with CNO had seizures. Interestingly, we discovered that we were not the first to have this happen. Alexander and colleagues (Alexander et al., 2009) report behavioral seizures in mice with Gq-coupled DREADDs (hM3Dq) in forebrain principal neurons with CNO doses as low as 0.5 mg/kg. Surprisingly, nearly 70% of animals that received a 5 mg/kg dose of CNO had seizures that lasted upwards of 120 min post-administration and sometimes even resulted in death. Due to these complications, we were not able to examine the role of IL/vmPFC terminals in the BLA in modulating anxiety-like behavior in this study. One of our interesting electrophysiology findings is that the glutamate release probabilities, measured using paired-pulse ratios, were significantly different between PL/dmPFC and IL/vmPFC terminals in the BLA under normal conditions (Figure 2). Specifically, we found that IL/vmPFC synapses had a higher release probability than PL/dmPFC synapses. A number of different mechanisms exist that could explain these differences in release probability including: size of the readily releasable pool, number of release sites, and alterations in calcium dynamics (Fioravante et al., 2011). Calcium plays an integral role in neurotransmitter release and is thought to underlie changes in short-term synaptic plasticity including release probability (reviewed in (Catterall et al., 2013; Neher and Sakaba, 2008; Südhof, 2012)). In order to examine the extent to which calcium played a role in determining glutamate release probability from PL/dmPFC and IL/vmPFC terminals in the BLA, we altered calcium levels in the external bath solution. We found that lowering extracellular calcium levels affected the release probability of IL/vmPFC, but not PL/dmPFC inputs suggesting that differences in calcium dynamics may underlie the differences in synaptic strength between these two circuits. Although our data demonstrate that calcium is a contributing factor, we do not know the exact mechanism underlying these differences. Some possibilities include differences in calcium influx, calcium-sensing proteins, calcium channel function, and calcium buffers. To our knowledge, no previous studies have directly compared the synaptic strength using paired-pulse ratios of these two pathways in rats. However, one study did report that these pathways recruited equivalent excitation and feedforward inhibition in BLA principal neurons (Arruda-Carvalho and Clem, 2014). It should be noted that their electrophysiology recordings were done in mice, and they never compared glutamate release probability of these synapses. One way to interpret our findings of differences in ‘basal’ glutamate transmission is that under ‘normal’ circumstances, the IL/vmPFC is exerting the primary top-down control of the BLA to prevent maladaptive behavior responses. However, during a pathological state, such as during alcohol withdrawal, the PL/vmPFC takes over at the primary sources of top-down control and drives maladaptive behaviors. Notably, alterations in synchronized activity between medial prefrontal and amygdala have been reported in human alcoholics (Müller-Oehring et al., 2015), although the behavioral implications of these changes were not directly assessed.

GABAergic transmission in the BLA serves an important role in providing both feedforward and feedback inhibition onto glutamatergic principal neurons to regulate the balance of inhibitory and excitatory transmission, which ultimately determines the output of the BLA. Our lab has examined the effects of chronic ethanol exposure and withdrawal on these GABAergic circuits and we found that withdrawal selectively decreases GABAergic transmission from feedforward neurons without altering the release from local interneurons (Diaz et al., 2011b). In the present study we examined the excitatory – inhibitory ratios of PL/dmPFC and IL/vmPFC synapses in the BLA (Figure 6) in both air-exposed control animals and those in withdrawal. In line with our paired-pulse findings, we found that the E/I ratios were differentially altered in these two pathways such that the E/I ratio was increased at PL/dmPFC – BLA synapses and decreased at IL/vmPFC – BLA synapses. These opposing effects were driven by a decrease in IPSC amplitude recruited from PL/dmPFC stimulation and an increase in IPSC amplitude resulting from IL/vmPFC stimulation in CIE/WD animals. Although we cannot say whether these GABAergic responses were feedforward or feedback in nature without recording from the GABAergic neurons themselves, we were able to demonstrate that these responses were disynaptic. This is evidences by significantly longer onset latencies for IPSC when compared to EPSCs. And, the IPSCs were not able to be rescued through the application of TTX and 4-AP suggesting they were action potential-dependent responses. Overall, these GABAergic data are demonstrating there is less recruitment of GABA from PL/vmPFC terminals, which makes sense in the context of increased presynaptic glutamate release; withdrawal from chronic ethanol exposure enhances glutamate release probability and decreases GABAergic recruitment in the PL/dmPFC pathway. Conversely, the findings that IL/vmPFC terminals are recruiting more GABA even though their glutamate release probability is decreased is harder to interpret. We believe that is represents one of the following scenarios; withdrawal from chronic ethanol exposure may cause the IL/vmPFC terminals to engage with a different population of neurons or the amount of stimulation we had to apply to these terminals to elicit comparable EPSCs in the AIR and CIE/WD animals may have been different, resulting in greater stimulation and GABAergic release in our withdrawal recordings. The latter hypothesis is feasible considering the fact that the prefrontal cortex has been shown the drive distinct populations of projection neurons in the BLA (Laura McGarry et al., 2017). In conclusion, our results indicate a novel role of discrete mPFC inputs to the BLA beyond that of fear. We demonstrate that the PL/dmPFC – BLA pathway is involved in regulating anxiety and that inhibiting these projections during alcohol withdrawal can decrease withdrawal-induced anxiety-like behavior. These data suggestion a mechanism by which withdrawal from chronic ethanol exposure selectively strengthens PL/dmPFC synapses while weakening IL/vmPFC synapses in the BLA. Strategies focused on reversing these alterations may offer targets for treatment development. In the future, it will be interesting to examine how the IL/vmPFC – BLA circuit modulates anxiety-like behavior as well as further parse apart the mechanisms underlying the ethanol-induced synaptic plasticity in these two circuits.

Author Contributions
MM and BM Designed Research; MM, BP, and AC Performed Research; MM Analyzed Data and Wrote the Paper.

